# A symbiont phage protein aids in eukaryote immune evasion

**DOI:** 10.1101/608950

**Authors:** M.T. Jahn, K. Arkhipova, S.M. Markert, C. Stigloher, T. Lachnit, L. Pita, A. Kupczok, M. Ribes, S.T. Stengel, P. Rosenstiel, B.E. Dutilh, U. Hentschel

## Abstract

Phages are increasingly recognized as important members of host associated microbial communities. While recent studies have revealed vast genomic diversity in the virosphere, the new frontier is to understand how newly discovered phages may affect higher order processes, such as in the context of host-microbe interactions. Here, we aim to understand the tripartite interplay between phages, bacterial symbionts and marine sponges. In a viromics approach, we discover 491 novel viral clusters and show that sponges, as filter-feeding organisms, are distinct viral niches. By using a nested sampling design, we show that each sponge individual of the four species investigated harbours its own unique virome, regardless of the tissue investigated. We further discover a novel, symbiont phage-encoded ankyrin domain-containing protein which appears to be widely spread in phages of many host-associated contexts including human. The ankyrin protein (ANKp) modulates the eukaryotic immune response against bacteria as confirmed in macrophage infection assays. We predict that the role of ANKp in nature is to facilitate co-existence in the tripartite interplay between phages, symbionts and sponges and possibly in many other host-microbe associations.

## Introduction

Phages are the most abundant and diverse entities in the oceans^1, 2^ and, along with their role as major bacterial killers, significantly impact global biochemical cycles^3^, bacterial fitness and diversity^4, 5^. In the last century, a plethora of fine-tuned defence, counter-defence mechanisms, and lysogenic conversion factors have been discovered through research focussing on phage-bacteria interactions^6, 7^. Importantly, however, in host-associated microbial communities, a third player, the eukaryotic host, not only sets the stage but may also interact with both other parties in its own interest. While recent sequencing-driven studies throughout animal taxa highlight phages as a ubiquitous part of host-associated microbiota, surprisingly little is known about the taxonomic association and mode of interaction between phages, their bacterial hosts and the animals that harbour the microbial communities^8^. While a direct phage-eukaryote interaction would be phagocytosis or transcytosis of virions by eukaryote cells, an indirect interaction would function via manipulation of the microbiome by the phage or eukaryote. The mechanisms of manipulation might consist of maintaining microbiome diversity by “killing the winner”^9^, maintaining its integrity by creating a barrier against invasion (the bacteriophage adherence to mucus [BAM] model^10^), or driving nutrient fluxes by the viral shunt^3^. However, evidence of phage-bacteria-eukaryote interplay is still scarce, and the underlying mechanisms are often unknown. Further, few studies systematically embraced complexity across biological scales ranging from protein- and cellular-level interactions up to viral association among tissues, individuals and species.

An attractive model that allows us to study host-microbe interactions in a natural environment are marine sponges, which are associated with stable, highly complex and specific microbial communities^11^. As filter-feeding animals, sponges pump up to 24,000 litres of seawater through their system per day^12^, exposing them to up to an estimated ∼2.4×10^13^ viruses daily. High exposure to phages, a major bacteriolytic element, raises questions about how microbiome homeostasis can be maintained. Morphologically, the lack of tissue boundaries along with the high bacterial cell densities would favour viral outbreaks within sponge tissues. Interestingly, defence mechanisms to alien nucleotides such as invading phages and plasmids are clearly enriched features of microbial sponge symbionts as indicated by single-cell genomics and metagenomics^13, 14, 15^. These defence mechanisms are based on self–non-self-discrimination (e.g., restriction-modification system) or prokaryotic adaptive immunity (i.e., CRISPR-Cas system), representing major strategies against viral infection. While these are indications for the selective advantage of phage resistance for the bacterial symbiont lifestyle, the beneficial effects of phages on the sponge microbial community are largely unexplored. The presence of virus-like particles in sponges was described early in 1978^16^ and was recently confirmed in an electron microscopy-based study counting 50 different viral morphotypes^17^. A first approach to sponge virome sequencing was established^18^ and applied to study viruses from Great Barrier Reef sponges collected from different locations and timepoints^19^. The comparison of the viral community profiles of the assayed species indicated species-specific viral signatures in sponges sharing low identity to known viral genomes. In the present study, we studied the tripartite phage-symbiont-eukaryote interplay by generating nested viromes from Mediterranean sponges and identifying sponge specialized taxa and host-enriched phage functions. We present the discovery of an immunomodulatory phage protein that has the potential to critically alter the interaction between sponges and microbial symbionts.

## Results

### High diversity and novelty in marine sponge viromes

We report the metagenomic analysis of marine sponge viromes that were sampled to cover the levels of sponge species, sponge individuals and sponge tissues. Viruses from nearby seawater, collected in immediate vicinity and at the same time, were used as controls. With 142Gbp of sequencing data from 32 sponges (two tissues × 4 individuals × 4 species) and 4 seawater reference viromes, this represents the deepest sequencing effort performed on sponge viruses to date. The final assembly contained 4,484 curated viral contigs representing population level genomes (>=5 kb, hereafter termed “BCvir” for viral populations of the North Western Mediterranean Coast close to Barcelona), representing 51.4% of all the read level data (Suppl. Table 1, Suppl. Fig. 1). Of these, 101 were circular with matching ends and represent putatively complete viral genomes. The remaining contigs (of which 1,649 were >=10 kb) were either putative linear genomes or genome fragments^20^. To investigate how the 4,484 BCvir populations were positioned in the known viral sequence space, we clustered our sequences with an extended sequence space of 11,901 viral genome sequences obtained from viral Refseq and the ActinophageDB as well as 29,922 assembled contigs from 130 publicly available marine viral communities. This analysis was based on shared gene content and detected 3,218 viral clusters (VCs, Fig. 1a). The 4,484 BCvir populations partitioned into 813 VCs (green) representing 25.3% (n=813 of 3,218 VCs) of the total viral diversity included in this extended database. Notably, most of the BCvir diversity consisted of viruses that were never detected before, as indicated by the fact that these VCs contained only BCvir contigs (n=491 of 3,218 VCs; 15.3%) (Fig. 1b), many of which shared no distant edge with other VCs. To ensure that this observation was not inflated by the shorter 5 kb contig length cut-off we initially applied to capture shorter ssDNA viruses, we performed the same analysis again using a more stringent 10 kb length filter as suggested in Roux, Emerson ^21^. With this approach, the 1,649 BCvir populations partitioned into 997 sponge VCs, of which 371 were novel (n=371 of 1,304 VCs; 28.5%) in the extended sequence space. VCs delineate approximately genus-level taxonomy in known viruses^22, 23^ with at least 371 sponge-derived VCs appear not to be part of the 803 viral genera currently listed by ICTV^24^ (via ViralRefseq; see complementary analyses Suppl. Note 1). Our virome dataset contained 3.9% BCvir populations that could be annotated at the family level, representing mainly bacteriophages of the Caudovirales families, Siphoviridae, Myoviridae, and Podoviridae (Suppl. Note 2, Suppl. Fig. 3). Rarefaction analysis indicated that more sponge viral diversity remains to be discovered as the curve has not reached saturation (Fig. 1c). These observations, combined with the limited taxonomic overlap with other marine environments (i.e., seawater, corals, sediment and sponges from Australia) (Fig. 1b), led us to the conclusion that sponges, even though constantly filtering seawater, represent distinct niches for viruses with potential for novel functions.

**Fig 1:**
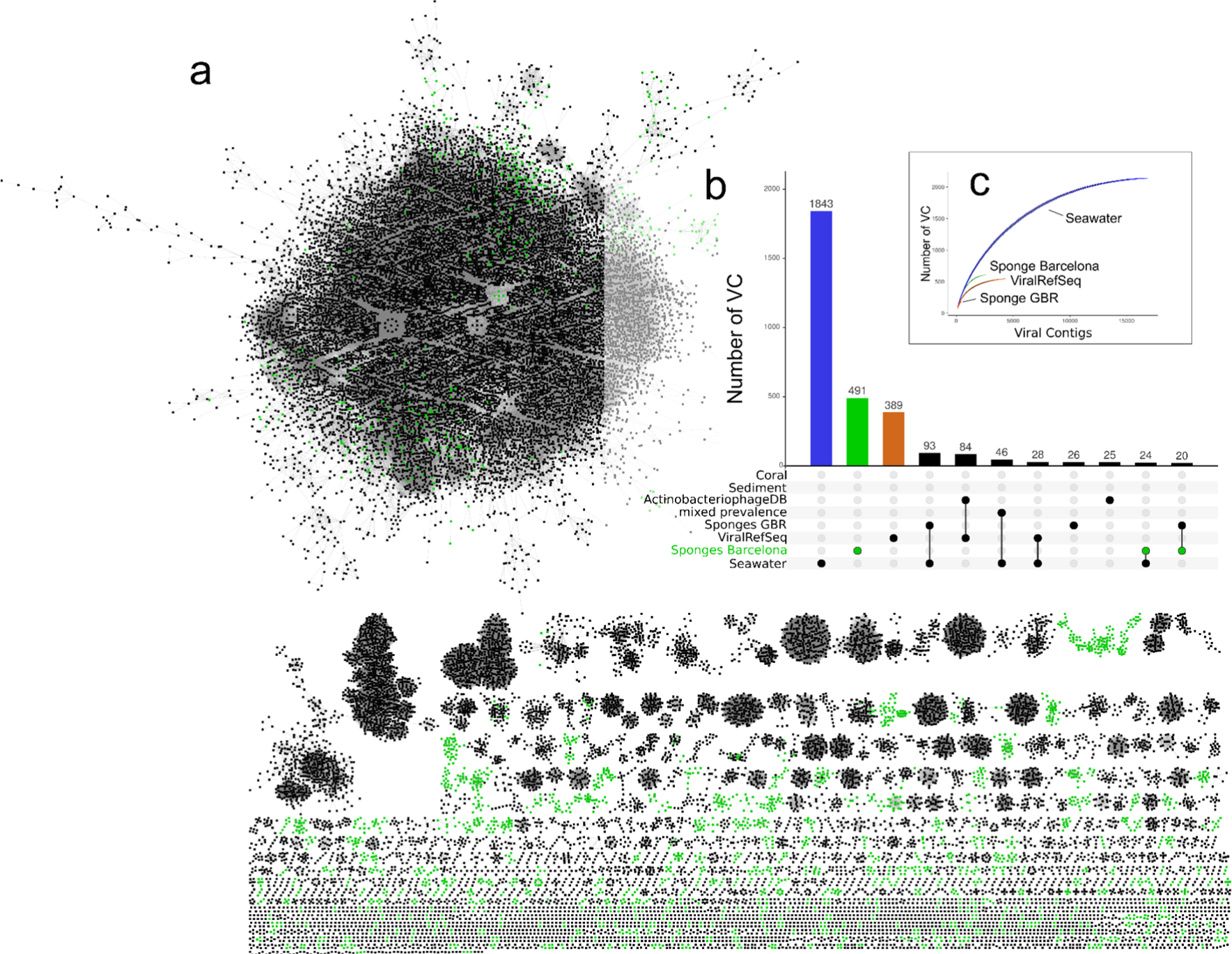
Sponge viruses in relation to known viral sequence space. (**a**) Nodes represent viral genomes or genome fragments and edges similarities based on gene content as estimated by the hypergeometric formula^22^. Sponge viral populations (BCvir, Sponge Barcelona), in green, were clustered with a custom database of 130 marine viromes, the ActinophageDB and RefSeqABVir, all in black. Viral clusters (VC; n=3,218 clusters) were identified by the Markov cluster (MCL) algorithm based on ICCC optimized inflation factor 1.4 (**b**) The matrix layout shows the number of VCs that are exclusive (one circle) or shared (multiple circles) between the eight different datasets used for clustering. Shown are the top intersections (>= 20 members) as vertical bar plot, sorted by size. (**c**) Rarefaction curves for the most diverse datasets showing the accumulation of VCs as a function of sampled viral contigs (N).

### Unique viromes in neighboring sponges

Viral communities of neighboring sponges were individually unique, host species specific and different from environmental seawater (Fig. 2 CrAss^25^ Clustering). This is indicated by the fact that viral community profiles grouped per sponge species (p value < 0.001, consistency value = 0.907) and were distinct from adjacent viroplankton (p value < 0.001, consistency value = 0.973). Variation in viral community composition within a given sponge species was mainly on the level of sponge individuals (p value < 0.001, consistency value = 0.679), rather than tissue specific signatures (p value = 0.991, consistency value = 0.534). These observations based on the fraction of cross contigs between the sample pairs (detailed in Methods) showed high concordance with results from hierarchical clustering of abundance profiles (Suppl. Fig. 7). We further explored viral populations by conceptualizing viral prevalence groups (see Methods for details). These were the generalists (prevalent in more than one sponge species or seawater), the specialists (prevalent in one sponge species or seawater), the individualists (detected in only one individual but both tissues) and intermediates (not falling into the above definitions) (Fig. 2). Notably, even though we obtained the samples from neighboring sponges of each sponge species at the same time point, individualists, at 38% (1,704 of 4,484 BCvir contigs), represented the largest virome group in our study. Furthermore, individualists were the second most abundant prevalence group in the dataset, indicating that these were not rare members of the community (Suppl. Fig. 8). In contrast, a minor fraction of BCvir population contigs were generalists being prevalent in all (n=10) or multiple (n=262) sample types (species/seawater). Specialists, with prevalence in one of the sponge species or seawater, were 27.9% (701 of 4,484). The intermediates, although not further categorized, still contain a species-specific pattern. Because efforts were made to minimize environmental variations by sampling in close spatial and temporal proximity, we conclude that the sponge individuals each have a unique viral fingerprint.

**Fig 2:**
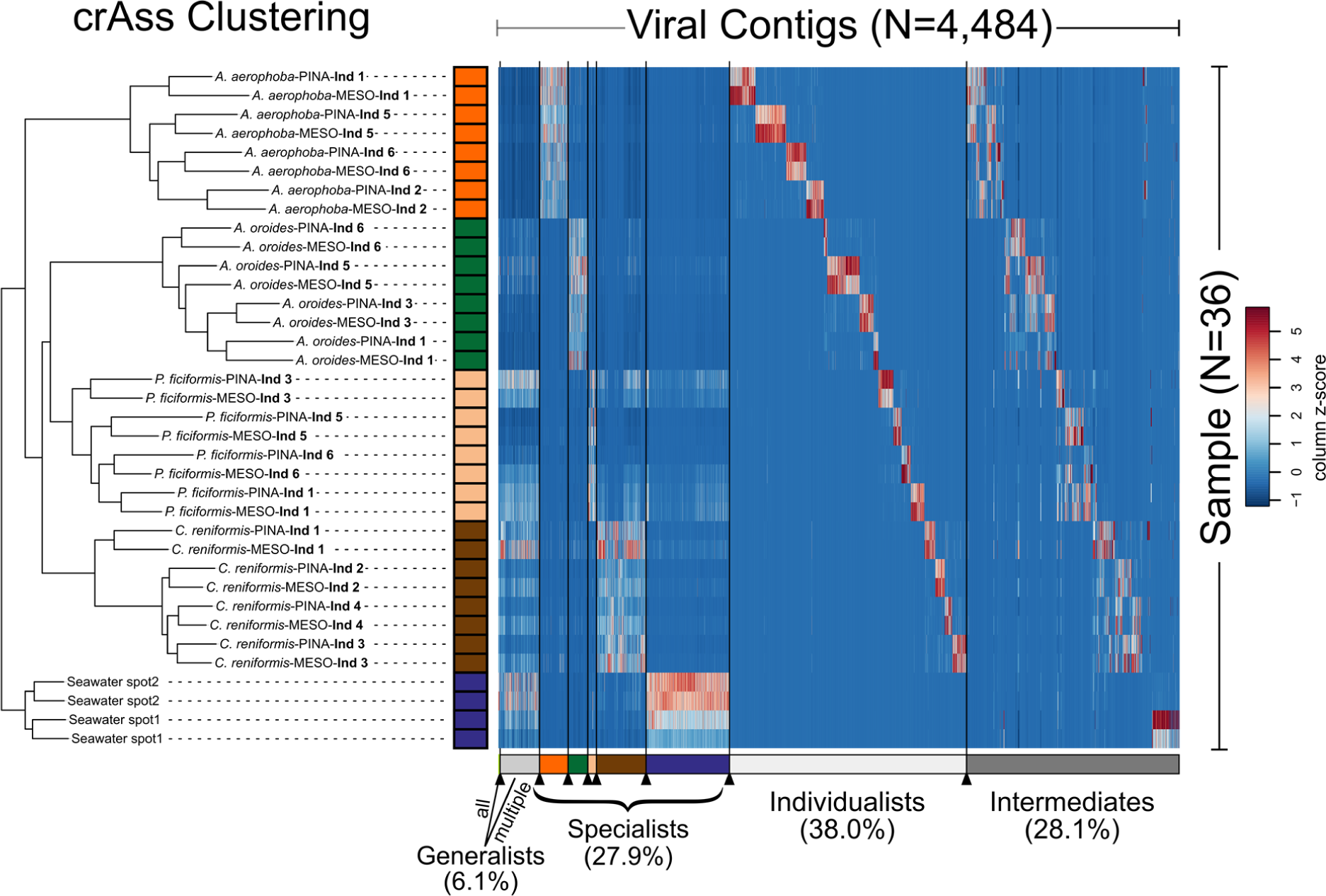
Iterative cross-assembly (crAss) of sponge-associated viromes. Clustering shows the distance between viral metagenomes based on the fraction of cross-assembled contigs between all sample pairs. Comparing topology against 1,000 random trees indicates significant separation of environments (sponge vs seawater) and sponge species (p value<=0.001) but not sponge tissues (p value=0.991). Heatmap shows the relative abundances of viral contigs and their grouping into prevalence groups as detailed in the Methods section. Color scheme is based on *z*-score distribution across samples from low (blue) to high (red).

### Symbiont phage protein aids bacteria in eukaryote immune evasion

To identify factors that might improve phages’ fitness in the interplay with their prokaryotic host cells and the extended eukaryotic host cells, we queried BCvir populations for auxiliary genes using a custom approach (see Methods for details). A first screening, as described in^26^, identified auxiliary metabolic genes (AMGs), which are detailed in Suppl. Data 1. Importantly, we then extended our search for cellular membrane transporters, adhesins, defence systems and cellular signal molecules owing to their potential relevance in a symbiosis context. We were surprised to find Ankyrin repeat domains (ANKs), discussed modulators of eukaryote-prokaryote interaction^27^, to be encoded on sponge-associated phages (Fig. 3a). These ANK encoding phages (BCvir 2964, BCvir 2161, BCvir 4986), which we will call Ankyphage 1, 2, 3 hereinafter (Suppl. Data 2; Ankyphage annotation), recruited reads from 12 of 32 sponge viromes but were absent in seawater. All three Ankyphages fall in the category “intermediates” (Fig. 2). Furthermore, Ankyphages were in the top 75^th^ percentile of most abundant viruses detected in *Aplysina* and *Chondrosia*. To ensure that Ankyphage sequences are indeed phage, we confirmed their phylogenetic placement among bacteriophages based on capsid alignments (see Suppl. Fig. 4) and ensured on the same contigs the presence of further phage domains, such as phage terminase (PF03354), phage portal protein (PF04860), and phage P22 coat protein (PF11651). The presence of *ANK* in CsCl-purified virus particles was confirmed by Sanger sequencing of the amplified *ANK* gene. Notably, the domain architecture of two Ankyphage ANKs comprised N-terminal signal peptides but no transmembrane domains (Fig. 3b). This suggests that these ANKs are secreted from a phage-infected prokaryotic cell (“virocell”) rather than that they function in the virion.

**Fig 3:**
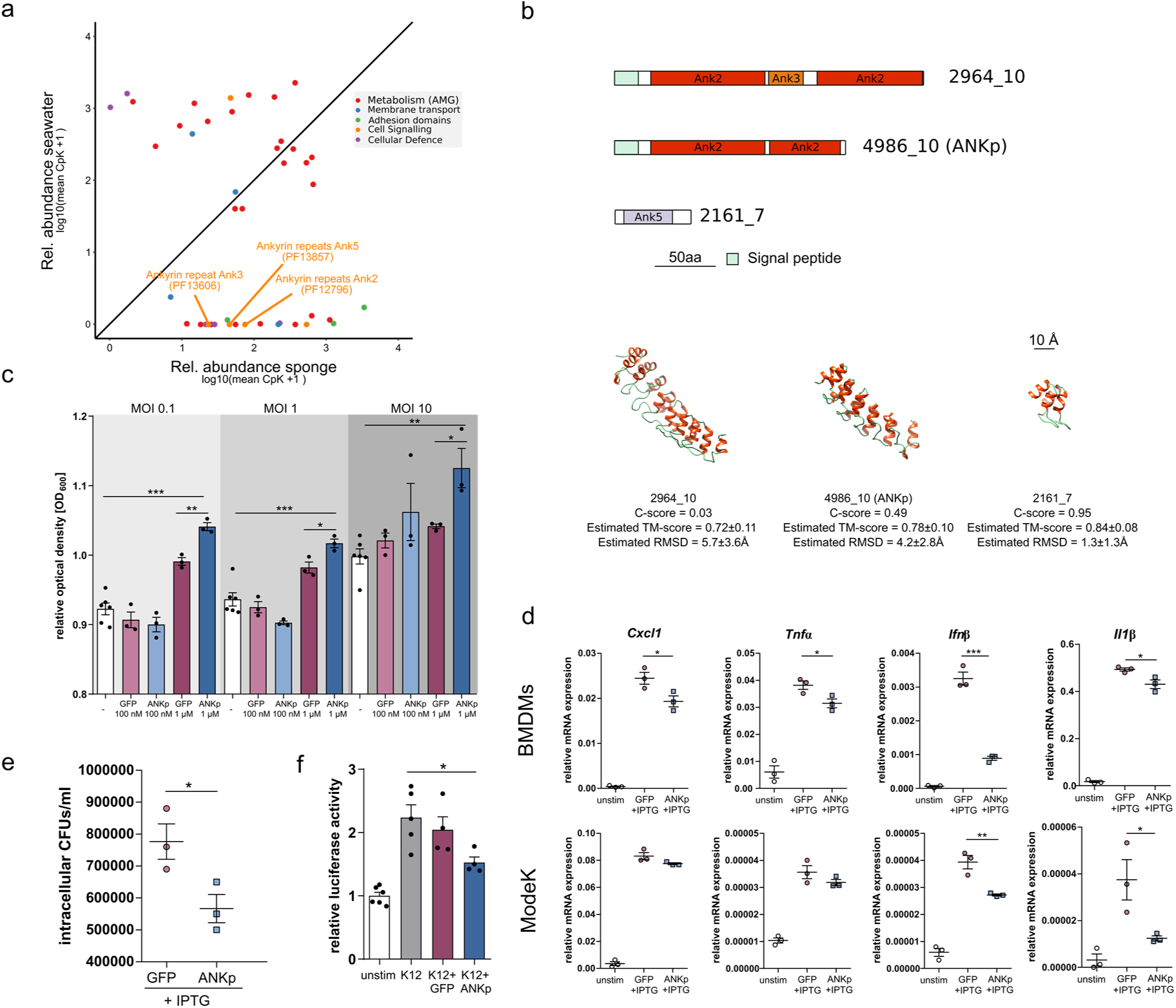
Symbiont phage ANKp reduces phagocytosis and immune response of eukaryote cells towards bacteria. (**a**) Scatterplot comparing relative abundance of auxiliary viral functions in sponge versus seawater (raw data and further host association factors available in Suppl. Data 1). (**b**) Domain architecture of ankyrin repeat encoding genes from sponge-enriched phages and representative protein models, approximated using I-TASSER. (**c**) Growth kinetics of *E. coli* K12 that are challenged with murine bone marrow-derived macrophages (BMDMs) upon preconditioning with GFP and ANKp at 100 nM or 1 µM. MOI refers to multiplicity of infection. The white bars indicate unstimulated (-) controls. Data are presented as the mean ± SEM of at least three independent experiments. (**d**) Expression levels of pro-inflammatory cytokines in macrophages (BMDMs) and intestinal epithelial cells ModeK upon infection with ANKp expressing *E. coli*. (**e**) Gentamycin protection assay reveals that ANKp expression leads to a reduced number of intracellular bacteria. (**f**) NF-□B expression in ModeK with dual-luciferase assay using an NF-□B– dependent firefly luciferase (pNF-□B-Luc; Clontech) and Renilla luciferase driven by a thymidine kinase promoter. Data are presented as the mean ± SEM of at least three independent experiments. Statistical significance between treatments was determined by two-tailed unpaired Student’s *t*-tests with *p < 0.05, **p < 0.01 and ***p < 0.001.

To examine whether phage-encoded ANKs can manipulate the bacteria-eukaryote interaction, we synthesized Ankyphage 3 ANK protein (ANKp) and assayed its impact on the interaction between bacteria and murine macrophages. Murine cell lines were chosen for the lack of an experimentally tractable model for sponge-microbe interactions^28^. When *E. coli* was pre-conditioned with purified ANKp, the bacterium survived in significantly higher abundances upon predatory pressure from murine macrophages compared to GFP controls (Fig. 3c, Suppl. Fig. 5). This effect was dose dependent (1 µM > 100 nM ANKp) and reproducible for different bacteria to macrophage ratios (multiplicity of infection (MOI) 0.1, 1 and 10). Gentamycin protection assays, which allowed us to quantify the intracellular bacteria fraction, showed that increased survival of ANKp pre-conditioned *E. coli* was paralleled with a decreased number of intracellular bacteria expressing the protein (p value 0.0414, t=2.963, df=4) (Fig. 3e). This indicates that ANKp-mediated bacterial survival is facilitated by decreased macrophage phagocytosis rates. To ensure that ANKp protein had no toxic effect on one of the players we performed bacterial growth experiments in culture and on plates and for the eukaryotic cell line MTS assays upon protein exposure showing no/low cytotoxicity (Suppl. Fig. 6). The same growth experiment exercised with pre-conditioned *Bacillus subtilis*, a Gram-positive representative, was consistent with ANKp-mediated bacterial survival during macrophage challenge (Suppl. Fig. 5).

On the side of the macrophage, ANKp synthesis by *E. coli* led to a reduced expression of pro-inflammatory cytokines upon bacterial exposure (Fig. 3d). Specifically, this included a reduction in tumour necrosis factor alpha (*T*NF-□), *C*xcl1, and *I*fn1β. To independently validate phage ANKp-mediated eukaryote immune suppression and to extend the analysis to a further eukaryotic cell type, we performed an NF-□B–dependent firefly luciferase assay on murine gut endothelial ModeK cells. In line with previous results, the NF-□B response, a central hub of eukaryote immunity^29^, was downregulated when ModeK cells were exposed to ANKp expressing *E. coli* (Fig. 3f). In summary, this shows that ANKp modulates the eukaryote response to bacteria by downregulating pro-inflammatory signalling along with reduced phagocytosis rates.

To investigate whether phage ankyrins are more common in nature, we extended homology searches to various other viral databases including IMGvr^30^ (July2018 release). We identified an abundant ANK encoding virus in the Great Barrier Reef sponge *Amphimedon queenslandica*^19^ (Suppl. Data 2; Ankyphage annotation). Furthermore, we identified ANKs in 418 predicted phage contigs deposited in IMGvr^30^ (Fig. 4). Notably, some, although not all, Ankyphages were obtained from host-associated environments such as the human oral cavity (n=67 phages), gut (n=6), stomach (n=5) or rhizosphere (n=4).

**Fig 4:**
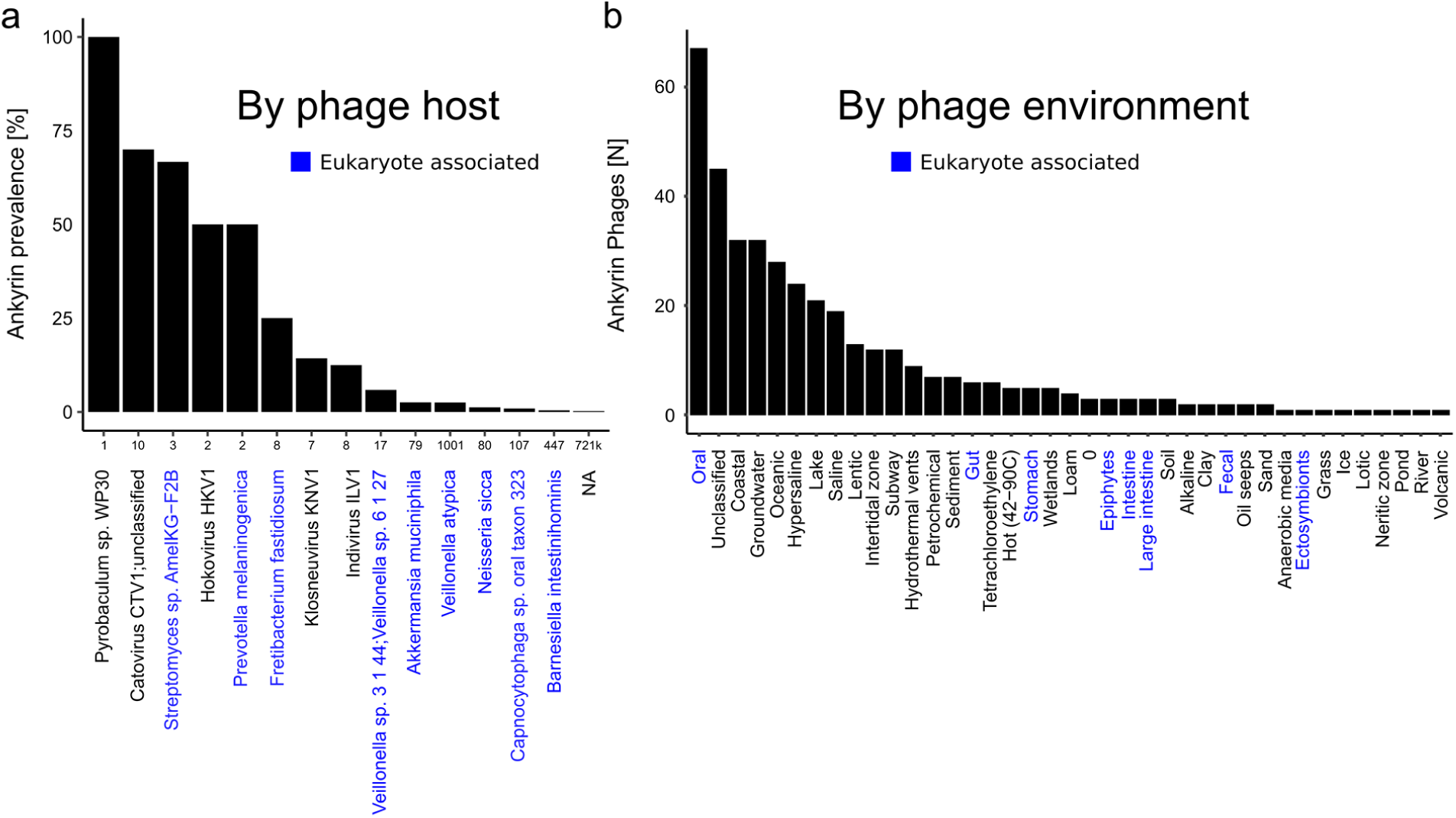
Global distribution of phage ANKs as deposited in the IMGvr database. (**a**) Prevalence of ankyrins in phages per predicted prokaryotic host (as deposited in IMGvr database metadata). Numbers on the x-axis indicate the total size of phage genomes/contigs per host taxon. (**b**) Number of phages with ANK domains per environment. Prokaryotes or environments with eukaryote host associations are highlighted in blue. The data are based on IMGvr database^30^ screening for Ankyrin Pfam signatures Ank (PF00023), Ank_2 (PF12796), Ank_3 (PF13606), Ank_4 (PF13637), and Ank_5 (PF13857) using InterPro (see Methods).

**Fig 5:**
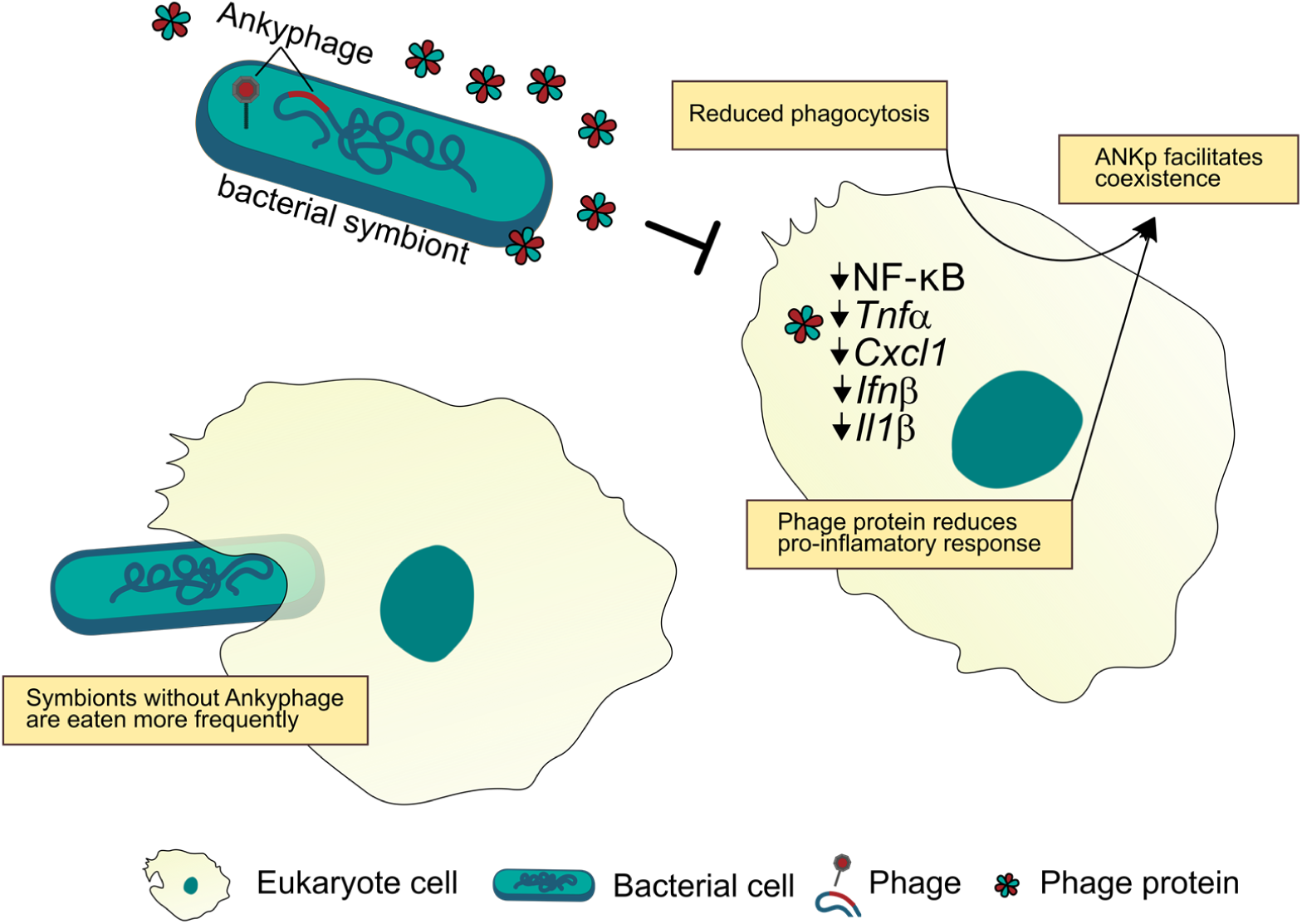
Proposed model for the phage-symbiont-eukaryote interplay. Ankyphages encode ANKp that, upon infection of the microbial symbiont, is expressed and secreted from the infected symbiont cell. When exposed to ANKp, eukaryote cells display a reduced inflammatory response to bacteria and eat less bacteria facilitating their coexistence.

## Discussion

In this study, we discovered a novel symbiont phage-encoded protein that modulates eukaryote-bacterium interaction by altering the eukaryotes’ physiology in response to bacteria. Furthermore, we followed an astonishingly intimate association between viruses and sponges across ecological scales (host tissue, individual, species) revealing unique individual fingerprints. This was achieved by an integrative approach where we combined tissue resolved virome sequencing on the scale of the sponge host community with synthesis-based cell assays.

We identified several hundred novel VCs in sponges that delineate genus-level taxonomy^22,23^. Notably, novelty was defined here by comparing to an extensive set of marine viral sequences, rendering it a rather conservative estimate. Although the applied genome-based network clustering approach cannot replace well-curated taxonomy^24^, the results of us and others^31^ indicate that marine animals indeed represent niches of distinct taxonomic viral diversity. Consequently, future efforts to capture more host-associated environments have high potential to add up to the increasingly understood planktonic virosphere of surface waters^26, 32^. Because animal microbiomes are highly species-specific^11, 33, 34^, and the virome depends on the microbiome *(and host)*, we further expect host-species-specific viral communities to rule in nature. This is supported by our sympatric sponge species, each holding characteristic viral communities (Fig. 3 clustering). Recently, Great Barrier Reef sponges were shown to contain similarly host specific viromes^19^. Systematic studies in other invertebrate systems such as *Hydra*^35^ and even signatures of phylosymbiosis in insects^36^ are likewise supportive for species-specific viromes associated with animals.

The next step was to resolve community-level signatures on the viral population scale (Fig. 2 heatmap) by integrating data of tissues and individuals in a temporally and spatially confined setting. This revealed that a considerable part of the sponge species-specific virome signatures was driven by viruses that were individually unique to sponge individuals and not to tissues (Fig. 2). Inter-individual differences were also the largest source of variance in the viromes of humans^37, 38^. To our knowledge, this report is the first that systematically extends this finding to marine animals. The high degree of individuality in sponge viromes was surprising considering the constant influx of nearby seawater that is filtered by sponges^39^, making it a conceivable route for horizontal transmission of phages among neighboring individuals. That this is not the case could either mean that transmission rates between nearby individuals are little or at least that rates do not keep pace with diversifying forces. These diversifying forces might be asynchronous temporal fluctuations between individuals, following delayed Lotka-Volterra-like dynamics^40^, or independent diversification from a source pool, as has been reported for other systems^41^. To resolve this, further studies capturing the temporal dynamics of environmental host-associated systems will be a valuable resource, because time adds an important dimension to viral diversity^42^. Furthermore, our results emphasize that small-scale collections of animals might easily underestimate true viral diversity by missing the abundant pool of individualist viruses present only on certain individuals.

The discovery of auxiliary ankyrin repeats (ANKs) in a new group of sponge-associated phages, we term Ankyphages, raised our special interest due to (a) their protein architecture, which indicates its secretion from the virocell, (b) their role as hubs of diverse protein-protein interactions, including functions in cellular signalling, and (c) their broader prevalence and abundance in the symbiotic context while lacking in nearby seawater. Taking these points and the fact that phages undergo strong pressure for genome reduction through purifying selection^43^, we reasoned that phage encoded ANKs might increase the fitness of the carrying phages in the context of the sponge holobiont. However, what is the phages’ advantage in orchestrating a virocell to secrete a protein binding domain? ANK repeats are widespread in all domains of cellular life^44^, but reports of ANKs in the world of phages are rare. A notable exception are PRANC-domains (Pox protein repeats of ankyrin CTD), which are ANK homologues of poxviruses being recently discovered in prophages of the ubiquitous insect endosymbiont Wolbachia^45^. Although the explicit function of these domains in the endosymbiont phage remains to be investigated, its placement in a conserved eukaryote association module is indicative of its functioning in the eukaryotes’ context^45^. In the bacterial world, ANKs were demonstrated to modulate interaction between species and even across kingdoms^46, 47^. A notable example is a previous study on sponges where chromosomally encoded ANKs from an uncultivated gammaproteobacterium seemed to modulate amoebal phagocytosis^27^ even though the underlying mechanisms on the eukaryote side are largely unclear. Inspired by these findings we estimated that phage-encoded ANKs might act to modulate bacteria-eukaryote interaction in favour of the virocell (Fig. 5).

Indeed, our *in vitro* experiments show that phage ANKp undermined eukaryote immune response towards bacteria and facilitated bacteria-eukaryote coexistence by reduced phagocytosis rates. To the best of our knowledge, this is the first secreted phage protein shown to downregulate eukaryote immune response and has important implications from a symbiotic perspective. The reduction of predatory pressure from the eukaryote host represents a selective advantage for in the symbiotic lifestyle of Ankyphage infected bacteria as compared to strains missing this trait (see Fig. 5). Eukaryote immune evasion by phage-mediated lysogenic conversion is an emerging field of research that is currently best studied in opportunistic pathogens^48^. Mechanisms range from phage-mediated reshaping of methicillin-resistant *Staphylococcus aureus* (MRSA) cell wall glycosylation to evade host immunity^49^, to yet to be determined genes of *Pseudomonas aeruginosa* phage Pf4 that downregulates eukaryote inflammatory response and at the same time is taming for non-invasive infection^50^. We are aware that the choice for the experimentally more approachable murine model can only be a proxy for processes yet to be observed in sponges. However, consistent signals in the phylogenetically distant system independent of bacterial Gram-state (*E. coli* and *B. subtilis*) and tested eukaryote cell type (macrophages, a critical component of innate immunity, and an epithelial cell line, representing an interface between microbes and hosts), might indicate a more widely distributed conserved mode of action. This is fuelled by our public database screenings where we found phage-encoded ANKs in other eukaryote-associated environments, such as phages inhabiting human cavities (oral, stomach, gut). In summary, our study highlights the novel diversity, intimate association, and tripartite interplay between phages, symbionts and the eukaryote host. Importantly, we identify and characterise a phage derived protein that can manipulate the immune interaction between eukaryotes and microbiota.

## Methods

### Nested sampling design

The high microbial abundance (HMA) sponge species *Petrosia ficiformis, Chondrosia reniformis*, and *Agelas oroides* (each n=4) were collected at the Mongrí Coast, Cala Foradada, 3°12’00.09’’E, 42°04’56.97’’N, Girona, Catalunya, Spain) close to Barcelona by snorkelling and scuba diving within a 20 m radius. We randomly sampled four individuals per species. *Aplysina aerophoba* (n=4) was collected at a different site 25 km south-west (see for metadata Suppl. Table 3). Immediately after collection, sponge samples were rinsed in sterile artificial seawater, plunge frozen in liquid nitrogen, and stored at −80°C until further processing. Prior to sponge sampling, for each spot, 30 litres of seawater were collected from the sponge vicinity using cooled sterilised tanks. Viroplankton was enriched by FeCl_3_ flocculation according to ^51^.

### Sample processing and virome sequencing

Deep frozen sponge individuals were dissected, separating the outer epithelial layer (pinacoderm) from the inner mesohyl matrix. All samples, including seawater references, were then randomly shuffled for virus purification and DNA/RNA extraction to avoid batch effects during processing. Samples were thawed in preboiled ice-cold extraction buffer and were disintegrated using a blender on ice at 6,500 rpm (T25 digital ULTRA-TURRAX, IKA). Particle aggregation was reduced by vortexing the suspension 10 min on ice. Tissue debris, PVPP bound secondary metabolites and bacterial cells were removed by centrifugation (2× 4,600g; 30 min at 4°C; ThermoScientific Heraeus Multifuge 3SR). The cleared supernatant was filtered through a 0.45 µm filter Conceicao-Neto, Zeller ^52^, and virions were pelleted using a Beckman SW-41-Ti swinging bucket rotor at 22,000 rpm for 2 h. Virions were re-suspended in SM-buffer overnight, purified by low speed centrifugation at 4,300g for 5 min, loaded onto a CsCl gradient (1.7/1.5/1.3/1.2/1.1) according Thurber, Haynes ^53^ and separated at 28,000 rpm for 2 h. Virion-containing layers (CsCl density 1.2-1.5) were retrieved using a syringe and confirmed for viral particles by epifluorescence and transmission electron microscopy. The virions were diluted in SM-buffer, purified by low speed centrifugation as before and pelleted at 28,000 rpm for 2 h. Upon overnight resuspension in Tris buffer, and removal of undissolved particles at 1000 g for 1 min, the supernatant was transferred to a fresh tube and incubated with benzonase for 2 h at 37°C to remove free nucleic acid contamination. Encapsulated viral DNA and RNA were extracted according to Thurber, Haynes ^53^ and ^54^. Viral nucleotides were randomly amplified using a modified version of the Whole Transcriptome Amplification Kit 2 (WTA2, Sigma Aldrich) as described in Conceicao-Neto, Zeller ^52^. This approach allows the capture of single-stranded and double-stranded DNA and RNA viruses with little amplification bias (as sequence reads represent viral genome copies). NexteraXT libraries were prepared and sequenced on a Hiseq2500 run with 2×250 bp paired end reads at IKMB Kiel (Suppl. Data 3). From the same tissue as used for the viromes, V1V2+V3V4 of the 16S rRNA gene was amplified and sequenced as described in Thomas, Moitinho-Silva ^11^.

### Metagenome cross-assembly and curation

Illumina reads were quality trimmed and cleaned from adapters, primers and reads with Ns or an average Q-score below 15 using BBMap v37.75 (https://sourceforge.net/projects/bbmap/). The reads were assembled together per library (n=36) and in random subsets of the total library pool (50× 0.01%, 12× 0.05%, 12 × 0.10%) using metaSPAdes v3.11.1 with default parameters. This multistep assembly strategy was tested in pilot assemblies to improve the quality of the assemblies as described in more detail in ^32^. Contigs from all assemblies were clustered using a custom script (https://github.com/kseniaarkhipova/RedRed) into populations with mummer3^55^ at >=95% ANI across >=80% of their lengths as inspired by ^21^. To filter for viral sequences and to remove remaining potential cellular contamination, population contigs were submitted to VirSorter 1.0.3 (using Virome Database and Virome decontamination options) and were additionally classified with the Contig Annotation Tool (CAT; https://github.com/dutilh/CAT). Contigs above 5 kb that were VirSorter classified and/or had superkingdom classification “Viruses” in CAT were used for downstream genome-centric analysis. For functional gene-centric analyses, we increased stringency against cellular sequence contamination by considering only contigs with at least two VirSorter hits for viral hallmark genes (i.e., “major capsid protein,” “portal”, “terminase large subunit,” “spike”, “tail,” “virion formation” or “coat) or CAT “viral superfamily” annotation. In the next round of cellular decontamination, we screened against single-copy prokaryotic marker genes using Anvi’o v.2.1.1 workflow^56^. A proportion of 6.19% (79 of 1276) of the contigs were hit by the single-copy prokaryotic marker database. Manual curation ensured that most of the hits were homologous to phage nucleotide replication machinery (DNA/RNA polymerases) and RecA, while there were no hits against any ribosomal RNA indicative of low remining contamination levels with cellular DNA/RNA^57^. One contig with the ClpX C4-type zinc finger domain was removed from further analysis due to its unclear viral association. These in silico filtration steps ensured that no cellular signals should have been included in the functional analysis.

### Gene content-based viral clustering

Evolutionary relationships between the viral genome (fragments) were inferred by implementing reticulate classification based on gene sharing as developed by Lima-Mendez, Van Helden ^22^. Briefly, we predicted 869,624 proteins from a set of 46,307 viral sequences (see details below) with PRODIGAL v2.6.3^58^ and detected pairwise similarities using all-by-all BLASTp, requiring a minimal bit score 50. Protein families were identified with the Markov cluster algorithm (MCL) using inflation factor 2 ^59^. All viral genomes were then compared to each other for shared protein family content, and the probability that similarity was by chance was estimated using a hypergeometric formula^22^. The resulting significance scores were corrected for multiple comparisons, and genome pairs with scores ≥ 0 were joined by an edge [see https://github.com/kseniaarkhipova/LMCLUST]. To define viral clusters (VCs) in the genome network, we determined 1.4 as the best MCL inflation factor based on ICCC (intracluster clustering coefficient) maximization as described in^23^. The curated viral contigs were clustered with well-characterized isolate genomes downloaded from the Actinobacteriophage database project (http://phagesdb.org/; January 2018) and ViralRefseq (January 2018). To investigate overlap with other marine environments, we also clustered with viral sequences from 130 environmental virome libraries. Specifically, 78 viromes cross-assembled from seawater (incl. Tara Oceans), corals and sediment^32^, 24 viromes from a seawater transect throughout the Mediterranean Sea^60^ and all viromes from sponges known to date supplemented with corals^19^. Viral clusters were taxonomically classified based on the placement of ViralRefseq entries in the network using a custom script [https://github.com/MartinTJahn/cluster_screener]. For each taxonomic rank, clusters were screened for Viral RefSeq entries and classified according to the supermajority (3/4) of their taxonomic annotations.

### Abundance profiles

Relative abundance patterns of viral genomes in the different samples were assessed by mapping quality control reads from each library against the curated genome-centric catalogue using BBMap 37.75 (option ambiguous=random, ANI >= 99%). The resulting 36 × 4484 count matrix was normalised for contig/virus length and library size to yield counts per kbp (CpK, Eq. (1)):

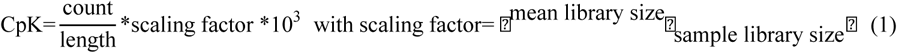

### Community patterns and prevalence group classification

Distances between viral metagenomes were computed with the reference-independent cross-assembly (crAss;^25^) tool. The resulting clustering was calculated based on the SHOT formula and was drawn with iTOL^61^. To infer significant clustering of sample categories (type, species, tissue), the leaf labels and associated sample categories were compared to N=1,000 trees where the leaf labels were randomized using a custom script [treestats.pl; https://github.com/linsalrob/crAssphage/tree/master/bin]. The observed patterns were validated independently by hierarchical clustering based on Bray-Curtis distances of community abundance signatures calculated in the R package vegan^62^.

Viral population enrichment for sample types was assessed using the population enrichment score (Eq. (2))

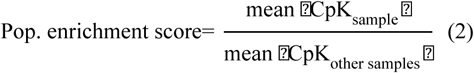

with a >= 2-fold enrichment considered to be enriched.

Viral population contigs (BCvir) were defined as detected in a sample when at least 75% of its length was covered by read mapping as suggested by ^21^. BCvir were considered prevalent based on the supermajority rule when detected in at least 75% of the samples in a sample category. “Individualists” were those BCvir that were detected in only one individual but both tissues. “Generalists” were prevalent in all or several sample categories, while “Specialists” were prevalent in only one sample type (Suppl. Data 4).

### Gene annotation and auxiliary gene classification

Proteins were predicted from viral contigs that passed the stringent cellular contamination filter (n=1,275 BCvir contigs) and were searched against the PFAM database (v31) using InterProScan v5.27-66.0^63^. Identified PFAM domains were then classified into 8 functional categories: “metabolism”, “lysis”, “structural”, “membrane transport, membrane-associated”, “DNA replication, recombination, repair, nucleotide metabolism”, “transcription, translation, protein synthesis”, “other”, and “unknown” as in ^64^ and extended by ^26^. This PFAM classification catalogue was augmented with manual classification of 85 PFAM signatures that were novel compared to the seawater viromes in the present study. We realized that in addition to auxiliary functions involved in the hosts metabolism (AMG), further categories might be relevant in the tripartite system of phage-prokaryote-eukaryote (PPE-interaction, hereafter). Therefore, we manually reannotated category “others” into classes “signalling and protein-protein interaction”, “cellular binding” and “cellular defence systems”, based on the literature research and functional evidence. The final extended PFAM classification catalogue for phages is given in Suppl. Data 5 and is open for further use in other systems. For abundance estimations, multiple identical PFAM motifs in one protein, such as by repeats, were counted as one to ensure that quantification is not biased towards repeat domains. Sequences were also annotated with Prokaryotic Virus Orthologous Groups (pVOGs; (Grazziotin *et al*. 2017)) through HMMER 3.1b2 (hmmscan -E 10^-5^), SEED subsystems through MG-RAST (E 10^-5^) and were matched to NCBI-nr database entries using Diamond (e-value 10^-5^ and identity >=40%). Proteins were screened for terminal signal peptides using SignalP v4.1f and for transmembrane domains using TMHMM 2.0c. The tertiary structure of phage ankyrin-containing proteins was approximated by I-TASSER (default settings)^65^. All annotations are combined in Suppl. Data 6.

### AnkP expression and purification

The 568 nt ANKp encoding phage gene (BCvir 4986 ORF 10) was optimized for *E. coli* codon usage and then de-novo synthesized in collaboration with GenScript (Piscataway, NJ, USA). The sequence was then cloned into the pET-22b(+) vector using BamHi and XhoI restriction sites. The same pET22b construct encoding GFP (Addgene plasmid # 38257 pET22b-GFP-CPDSalI) was used as a negative control and was a gift from Matthew Bogyo & Aimee Shen ^66^. Plasmids were transformed into BL21(DE3)-competent *E. coli*. Heterologous protein expression was induced in *E. coli* with 0.4 mM IPTG, and cultures were grown for 3 h at 37°C. For the native purification of the His-tagged target proteins, the Ni-NTA Fast Start Kit (Qiagen) was applied according to the manufacturer’s instructions.

### Cell assays

#### Murine bone marrow-derived macrophages (BMDMs)

C57BL/6 mice (10-14 weeks old) were killed, femur and tibia were dissected, and bone marrow was isolated under sterile conditions. Haematopoietic stem cells differentiated into BMDMs by incubating for 7 days in BMDM medium (1:1 SFM:DMEM, Gibco) supplemented with 10 % FCS (Biochrom), 1 % penicillin/streptomycin (Gibco), 1 % amphotericin B (Gibco), and 20 ng/mL macrophage colony-stimulating factor (mCSF, Immunotools). All animal experiments were approved by the Animal Investigation Committee of the University Hospital Schleswig-Holstein (Campus Kiel, Germany; acceptance no.: V242-7224.121-33) and were performed according the relevant guidelines and regulations.

#### Growth dynamics of conditioned *E. coli* and *B. subtilis* cultures

*E. coli* K12 (DSMZ, #498) or *B. subtilis* (NCIB 3610) overnight Luria–Bertani (LB) medium cultures were harvested at 4,000g for 10□min at room temperature and were washed twice with PBS. The bacterial suspension was then either conditioned with ANKp, GFP or PBS for 10 min at 4°C under mild agitation. Prior to infection, the medium of BMDM cell culture was replaced with fresh antibiotic-free BMDM medium, and the cells were then infected with conditioned *E. coli* K12 (0 nM, 100 nM, 1 µM purified protein) using a range of multiplicities of infection (MOIs). Optical density measurements at a wavelength of 600 nm were performed using a Tecan Infinite 200 plate reader in a 96-well plate as described in ^67^.

#### RNA extraction and quantitative RealTime PCR

Total RNA was isolated from BMDM and ModeK cell culture 16 h post infection using the RNeasy kit (Qiagen) and reverse transcribed using the Maxima H Minus First Strand cDNA Synthesis kit (Thermo Scientific). Quantitative RealTime PCRs were performed with TaqMan Gene Expression Master Mix (Applied Biosystems) according to the manufacturer’s instructions and were analysed on the 7900HT Fast Real Time PCR System (Applied Biosystems). The applied TaqMan assays for pro-inflammatory markers are listed Suppl. Table 4).

#### NF-κB–dependent luciferase assay

The dual-luciferase assay using an NF-□B–dependent firefly luciferase (pNF-□B-Luc; Clontech) and a Renilla luciferase driven by the thymidine kinase promoter (pRLTK; Clontech) was performed according to the manufacturer’s instructions. Briefly, ModeK cells cultured in DMEM (DMEM Glutamax plus 10% FCS, non-essential amino acids and HEPES, Gibco) were transfected with 20 ng pNF-□B-Luc and 3 ng pRL-TK using FuGENE 6 (Roche). Transfected cells were incubated for 24 h (37°C, 5% CO_2_), lysed and the lysate was subjected to the dual-luciferase assay carried out on a Tecan 96-well microplate reader.

## Data availability

All sequencing libraries and the cross-assembly, of both the sponge viromes and seawater references, as well as microbial amplicon data, are deposited in NCBI SRA database (NCBI BioProject accession: PRJNA522695). All custom code is available at GitHub as indicated in the methods section. ⍰

## Acknowledgements

We acknowledge funding by the DFG CRC1182 to U.H (TPB1), T.L (TPA4), P.R (TPC2), A.K. (TPC3). M.T.J. was supported by a grant of the German Excellence Initiative to the Graduate School of Life Sciences, University of Wuerzburg, and a Young Investigator Award of the CRC1182. S.M.M was supported by the Studienstiftung des Deutschen Volkes (German National Academic Foundation). We thank Laura Rix (GEOMAR), Rafel Coma (CEAB-CSIC) and Berta Pintó for sponge sampling.

## Contributions

M.T.J designed and performed the experiments, analysed the data, prepared figures and tables, and wrote the paper; L.P and M.R were involved in the planning and execution of sponge field sampling; T.L purified viral particles and extracted nucleotides; S.T.S and P.R performed cell culture experiments and immuno assays; M.T.J and S.M.M performed microscopy under supervision of C.S; B.E.D, A.K and K:A advised bioinformatic analyses and commented on analysis strategy; U.H helped in experimental design, data interpretation and reviewed drafts of the paper; All authors edited and approved the final version of the manuscript.

## Bibliography

1. Wommack KE, Colwell RR. Virioplankton: viruses in aquatic ecosystems. Microbiol Mol Biol Rev 64, 69–114 (2000).

2. Rohwer F. Global phage diversity. Cell 113, 141 (2003).

3. Suttle CA. Marine viruses--major players in the global ecosystem. Nat Rev Microbiol 5, 801–812 (2007).

4. Betts A, Gray C, Zelek M, MacLean RC, King KC. High parasite diversity accelerates host adaptation and diversification. Science 360, 907–911 (2018).

5. Marston MF, et al. Rapid diversification of coevolving marine *Synechococcus* and a virus. Proc Natl Acad Sci USA 109, 4544–4549 (2012).

6. Barrangou R, et al. CRISPR provides acquired resistance against viruses in prokaryotes. Science 315, 1709–1712 (2007).

7. Kronheim S, et al. A chemical defence against phage infection. Nature 564, 283–286 (2018).

8. Keen EC, Dantas G. Close encounters of three kinds: bacteriophages, commensal bacteria, and host immunity. Trends Microbiol 26, 943–954 (2018).

9. Rodriguez-Valera F, et al. Explaining microbial population genomics through phage predation. Nat Rev Microbiol 7, 828–836 (2009).

10. Barr JJ, et al. Bacteriophage adhering to mucus provide a non-host-derived immunity. Proc Natl Acad Sci USA 110, 10771–10776 (2013).

11. Thomas T, et al. Diversity, structure and convergent evolution of the global sponge microbiome. Nat Commun 7, 11870 (2016).

12. Weisz JB, Lindquist N, Martens CS. Do associated microbial abundances impact marine demosponge pumping rates and tissue densities? Oecologia 155, 367–376 (2008).

13. Horn H, et al. An enrichment of CRISPR and other defense-related features in marine sponge-associated microbial metagenomes. Front Microbiol 7, 1751 (2016).

14. Podell S, et al. Pangenomic comparison of globally distributed Poribacteria associated with sponge hosts and marine particles. ISME J, (2018).

15. Slaby BM, Hackl T, Horn H, Bayer K, Hentschel U. Metagenomic binning of a marine sponge microbiome reveals unity in defense but metabolic specialization. ISME J 11, 2465 (2017).

16. Vacelet J, Gallissian M-F. Virus-like particles in cells of the sponge Verongia cavernicola (Demospongiae, Dictyoceratida) and accompanying tissues changes. J Invertebr Pathol 31, 246–254 (1978).

17. Pascelli C, Laffy PW, Kupresanin M, Ravasi T, Webster NS. Morphological characterization of virus-like particles in coral reef sponges. PeerJ 6, e5625–e5625 (2018).

18. Laffy PW, et al. HoloVir: a workflow for investigating the diversity and function of viruses in invertebrate holobionts. Front Microbiol 7, (2016).

19. Laffy PW, et al. Reef invertebrate viromics: diversity, host specificity and functional capacity. Environ Microbiol Epub ahead of print, (2018).

20. Roux S, et al. Minimum Information about an Uncultivated Virus Genome (MIUViG). Nat Biotechnol, (2018).

21. Roux S, Emerson JB, Eloe-Fadrosh EA, Sullivan MB. Benchmarking viromics: an in silico evaluation of metagenome-enabled estimates of viral community composition and diversity. PeerJ 5, e3817 (2017).

22. Lima-Mendez G, Van Helden J, Toussaint A, Leplae R. Reticulate representation of evolutionary and functional relationships between phage genomes. Mol Biol Evol 25, 762–777 (2008).

23. Roux S, Hallam SJ, Woyke T, Sullivan MB. Viral dark matter and virus—host interactions resolved from publicly available microbial genomes. Elife 4, (2015).

24. King AMQ, et al. Changes to taxonomy and the International code of virus classification and nomenclature ratified by the international committee on taxonomy of viruses (2018). Arch Virol, (2018).

25. Dutilh BE, et al. Reference-independent comparative metagenomics using cross-assembly: crAss. Bioinformatics 28, 3225–3231 (2012).

26. Roux S, et al. Ecogenomics and potential biogeochemical impacts of globally abundant ocean viruses. Nature 537, 689 (2016).

27. Nguyen MT, Liu M, Thomas T. Ankyrin-repeat proteins from sponge symbionts modulate amoebal phagocytosis. Mol Ecol 23, 1635–1645 (2014).

28. Pita L, Fraune S, Hentschel U. Emerging Sponge Models of Animal-Microbe Symbioses. Front Microbiol 7, (2016).

29. Li Q, Verma IM. NF-kappaB regulation in the immune system. Nat Rev Immunol 2, 725–734 (2002).

30. Paez-Espino D, et al. IMG/VR v.2.0: an integrated data management and analysis system for cultivated and environmental viral genomes. Nucleic Acids Res 47, D678–d686 (2019).

31. Paez-Espino D, et al. Uncovering earth’s virome. Nature 536, 425–430 (2016).

32. Coutinho FH, et al. Marine viruses discovered via metagenomics shed light on viral strategies throughout the oceans. Nat Commun 8, 15955 (2017).

33. Moeller AH, et al. Cospeciation of gut microbiota with hominids. Science 353, 380–382 (2016).

34. Hacquard S, et al. Microbiota and host nutrition across plant and animal kingdoms. Cell Host Microbe 17, 603–616 (2015).

35. Grasis JA, et al. Species-specific viromes in the ancestral holobiont *Hydra*. PLoS One 9, e109952 (2014).

36. Leigh BA, Bordenstein SR, Brooks AW, Mikaelyan A, Bordenstein SR. Finer-scale phylosymbiosis: insights from insect viromes. mSystems 3, e00131–00118 (2018).

37. Abeles SR, et al. Human oral viruses are personal, persistent and gender-consistent. ISME J 8, 1753–1767 (2014).

38. Moreno-Gallego JL, et al. Virome diversity correlates with intestinal microbiome diversity in adult monozygotic twins. Cell Host & Microbe 25, 261-272.e265 (2019).

39. Taylor MW, et al. ‘Sponge-specific’ bacteria are widespread (but rare) in diverse marine environments. ISME J 7, 438–443 (2013).

40. Parsons RJ, Breitbart M, Lomas MW, Carlson CA. Ocean time-series reveals recurring seasonal patterns of virioplankton dynamics in the northwestern Sargasso Sea. ISME J 6, 273–284 (2012).

41. Enav H, Kirzner S, Lindell D, Mandel-Gutfreund Y, Beja O. Adaptation to sub-optimal hosts is a driver of viral diversification in the ocean. Nat Commun 9, 4698 (2018).

42. Arkhipova K, et al. Temporal dynamics of uncultured viruses: a new dimension in viral diversity. ISME J, (2017).

43. Duffy S, Shackelton LA, Holmes EC. Rates of evolutionary change in viruses: patterns and determinants. Nat Rev Genet 9, 267–276 (2008).

44. Jernigan KK, Bordenstein SR. Ankyrin domains across the tree of life. PeerJ 2, e264 (2014).

45. Bordenstein SR, Bordenstein SR. Eukaryotic association module in phage WO genomes from *Wolbachia*. Nat Commun 7, 13155 (2016).

46. Lambert C, et al. Ankyrin-mediated self-protection during cell invasion by the bacterial predator *Bdellovibrio bacteriovorus*. Nat Commun 6, 8884 (2015).

47. Wong K, Perpich JD, Kozlov G, Cygler M, Abu Kwaik Y, Gehring K. structural mimicry by a bacterial F Box effector hijacks the host ubiquitin-proteasome system. Structure (London, England: 1993) 25, 376–383 (2017).

48. Van Belleghem J, Dabrowska K, Vaneechoutte M, Barr J, Bollyky P. Interactions between bacteriophage, bacteria, and the mammalian immune system. Viruses 11, 10 (2018).

49. Gerlach D, et al. Methicillin-resistant *Staphylococcus aureus* alters cell wall glycosylation to evade immunity. Nature 563, 705–709 (2018).

50. Secor PR, et al. Filamentous bacteriophage produced by *Pseudomonas aeruginosa* alters the inflammatory response and promotes noninvasive infection *in vivo*. Infect Immun 85, e00648–00616 (2016).

51. John SG, et al. A simple and efficient method for concentration of ocean viruses by chemical flocculation. Environ Microbiol Rep 3, 195–202 (2011).

52. Conceicao-Neto N, et al. Modular approach to customise sample preparation procedures for viral metagenomics: a reproducible protocol for virome analysis. Sci Rep 5, 16532 (2015).

53. Thurber RV, Haynes M, Breitbart M, Wegley L, Rohwer F. Laboratory procedures to generate viral metagenomes. Nat Protoc 4, 470–483 (2009).

54. Lachnit T, Thomas T, Steinberg P. Expanding our understanding of the seaweed holobiont: RNA viruses of the red alga *Delisea pulchra*. Front Microbiol 6, 1489–1489 (2016).

55. Kurtz S, et al. Versatile and open software for comparing large genomes. Genome Biol 5, R12 (2004).

56. Eren AM, et al. Anvi’o: an advanced analysis and visualization platform for ‘omics data. PeerJ 3, e1319 (2015).

57. Roux S, Krupovic M, Debroas D, Forterre P, Enault F. Assessment of viral community functional potential from viral metagenomes may be hampered by contamination with cellular sequences. Open Biol 3, (2013).

58. Hyatt D, Chen G-L, Locascio PF, Land ML, Larimer FW, Hauser LJ. Prodigal: prokaryotic gene recognition and translation initiation site identification. BMC Bioinformatics 11, (2010).

59. Enright AJ, Dongen S, Ouzounis CA. An efficient algorithm for large-scale detection of protein families. Nucleic Acids Res 30, (2002).

60. López-Pérez M, Haro-Moreno JM, Gonzalez-Serrano R, Parras-Moltó M, Rodriguez-Valera F. Genome diversity of marine phages recovered from Mediterranean metagenomes: Size matters. PLoS Genet 13, e1007018 (2017).

61. Letunic I, Bork P. Interactive Tree Of Life (iTOL): an online tool for phylogenetic tree display and annotation. Bioinformatics 23, 127–128 (2007).

62. Dixon P. VEGAN, a package of R functions for community ecology. J Veg Sci 14, 927–930 (2003).

63. Jones P, et al. InterProScan 5: genome-scale protein function classification. Bioinformatics 30, 1236–1240 (2014).

64. Hurwitz BL, Brum JR, Sullivan MB. Depth-stratified functional and taxonomic niche specialization in the ‘core’ and ‘flexible’ pacific ocean virome. ISME J 9, 472–484 (2015).

65. Roy A, Kucukural A, Zhang Y. I-TASSER: a unified platform for automated protein structure and function prediction. Nat Protoc 5, 725–738 (2010).

66. Shen A, et al. Simplified, enhanced protein purification using an inducible, autoprocessing enzyme tag. PLoS One 4, e8119 (2009).

67. Erez Z, et al. Communication between viruses guides lysis–lysogeny decisions. Nature 541, 488–493 (2017).

