## Supplementary Figures and Tables for "A symbiont phage protein aids in eukaryote immune evasion"

Jahn *et al*.

**Supplementary Figures**

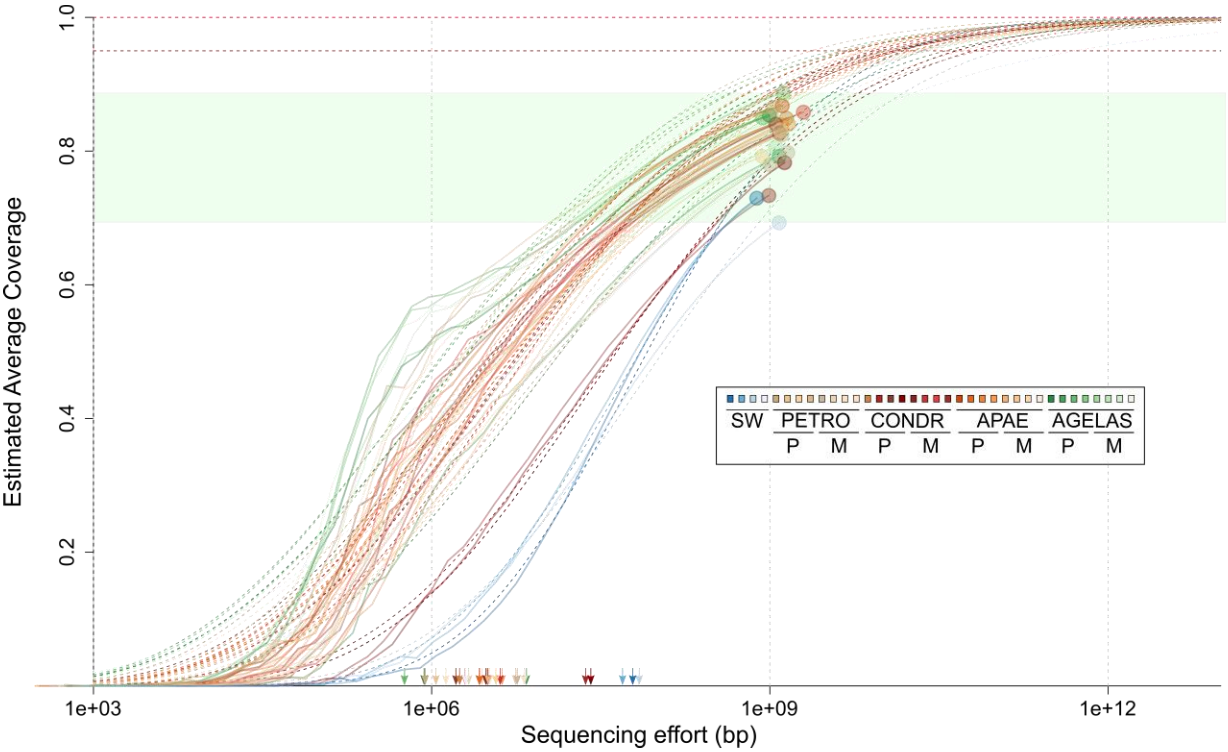


**Supplementary Figure 1: Community coverage of each metagenomic dataset was estimated using Nonpareil v3.301 with default parameters[1]**. The dashed lines indicate the fitted models of the Nonpareil curves (solid). Coloured circles are the estimated coverage for each sample, and the green area is the observed range for the samples. Horizontal dashed lines indicate 100 and 95% coverage. Coloured arrows indicate the needed sequencing effort to reach 50% coverage of the fitted model as a proxy for detected diversity. Abbreviations: SW = seawater, PETRO = *Petrosia*, CONDR = *Chondrosia*, APAE = *Aplysina*, AGELAS = *Agelas*, M = mesohyl, P = pinacoderm.

**Suppl. Note 1:** Genome-based network cluster validations

To ensure that our approach of defining viral clusters (VCs) grouped viral genomes at the genus level, we confirmed for ViralRefseq entries that there were (a) few clusters per genus and (b) few genera per cluster. Most ViralRefseq entries clustered in one cluster per given genus (Suppl. Fig. 2; 183 of 215 genera). The strongest exception was the genus Begomovirus, where genomes were distributed into 30 adjacent viral clusters. This is in line with reports stressing the need for new taxonomic demarcation criteria in this genus[2]. Multiple genera per cluster were observed in 1.8% (59 of 3,218) of the VCs in the applied network, representing a small minority of cases where our approach may have been too inclusive.

.

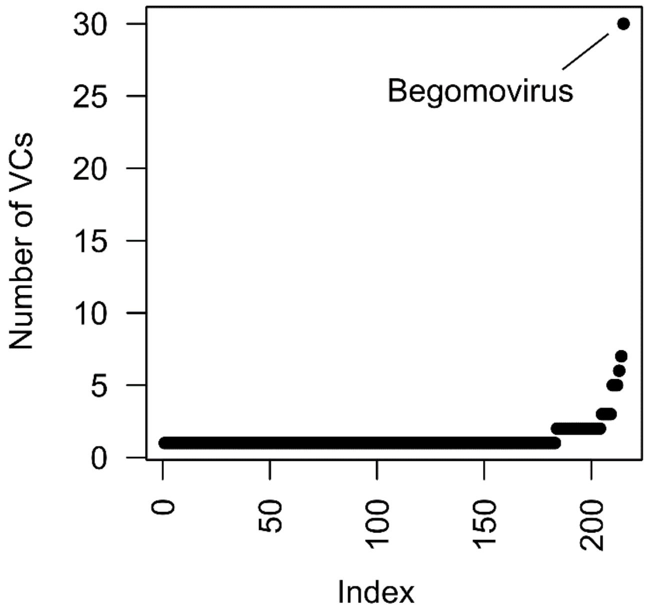


**Supplementary Figure 2:** Assigned viral clusters (VCs) per ViralRefseq genus. The index represents the 215 viral genera used for classification, where the number of assigned VCs per genus is 1, where VC delineation represents described genera.

**Suppl. Note 2:** Taxonomic affiliation of BCVir populations

The co-clustering with ViralRefseq representatives of known taxa allowed us to classify 1.2% (55/4,484) of BCvir populations on the genus level and 3.9% (177/4,484) on the family level. The predominant viral families where taxonomy could be applied represented single- and double-stranded DNA bacteriophages. These were tailed bacteriophages of the order Caudovirales (Podo-, Myo-, Siphoviridae) and members of the Microviridae subfamily Gokushovirinae (Suppl. Fig. 3). We further note the detection of ssDNA(+/-) Ambidensovirus, eukaryote viruses of the family Parvoviridae, in a majority of sponge individuals, which is consistent with observations from Australian reef sponges[3].

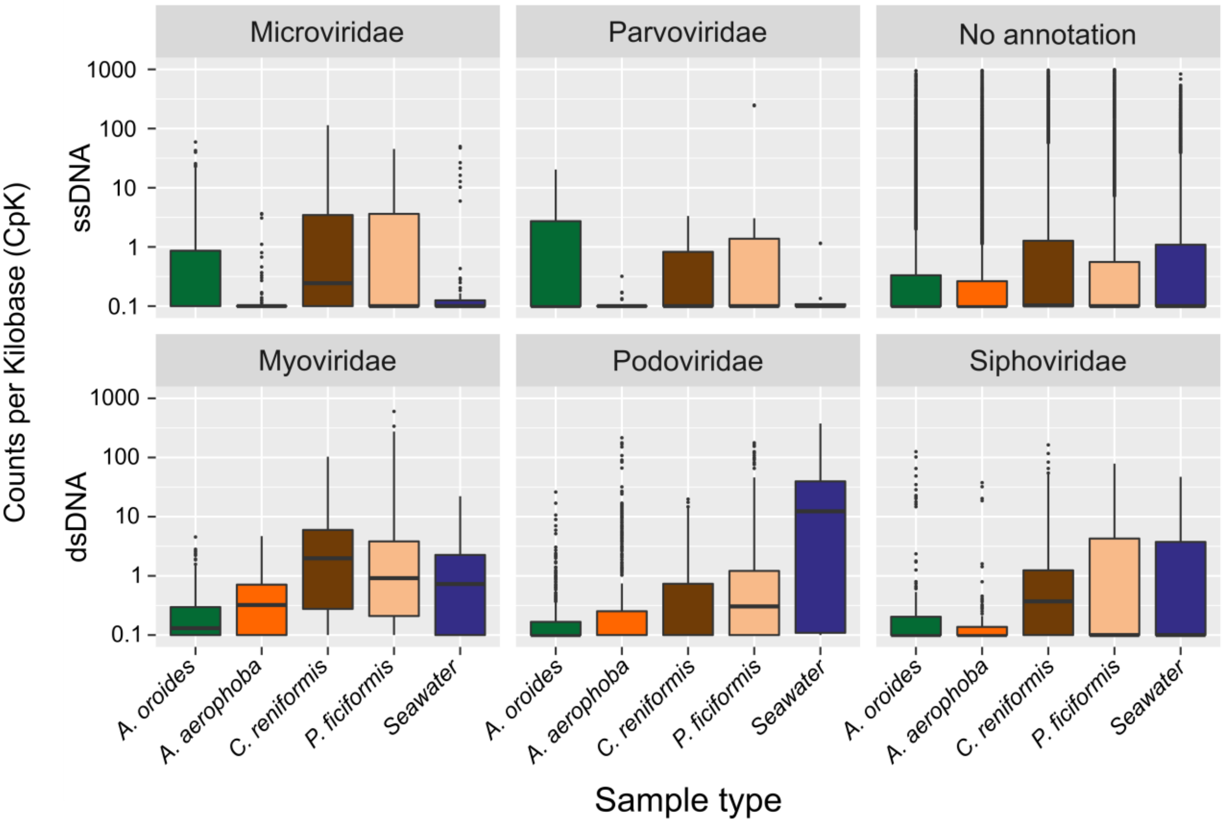


**Supplementary Figure 3: Taxonomic affiliation of viral population contigs from sponges.** This was based on clustering of BCvir populations with RefSeqABVir using 75% majority rule on family level. Overall, 3.9% (59 of 4,484) of the population contigs were assigned to known sequences on the family level.

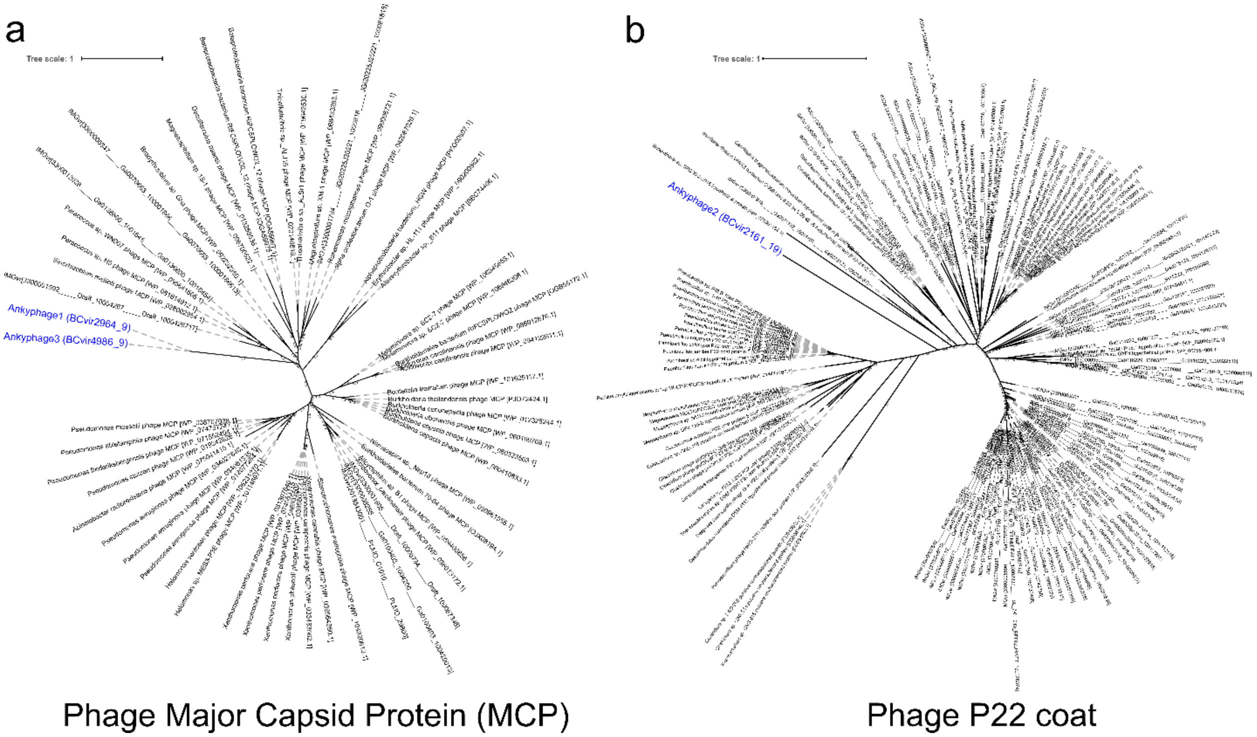


**Supplementary Figure 4: Phylogenetic analysis of structural genes places Ankyphages among bacteriophages.** Phylogenies were constructed based on (**a**) 401 aa of Phage major capsid protein (MCP) and (**b**) 382 aa of Phage P22 coat proteins with IQ-TREE[4] (1,000 bootstraps). The protein set to compare with was identified by BLASTx searches against GenBank (v. April2018) and IMGVr (v. July2018) (e-value ≤ $10^{-5}$; coverage ≥ 33% MVC and 25% P22). The set was deduplicated with usearch v8.1.1861 at 100% ANI, aligned with MUSCLE, and alignment was curated using Guidance2 ([5], column score >= 0.6). The best-fit model of protein evolution (LG+I+G+F for MVC, LG+R6 for P22) was identified with IQ-TREE. Trees were plotted using iTOL.

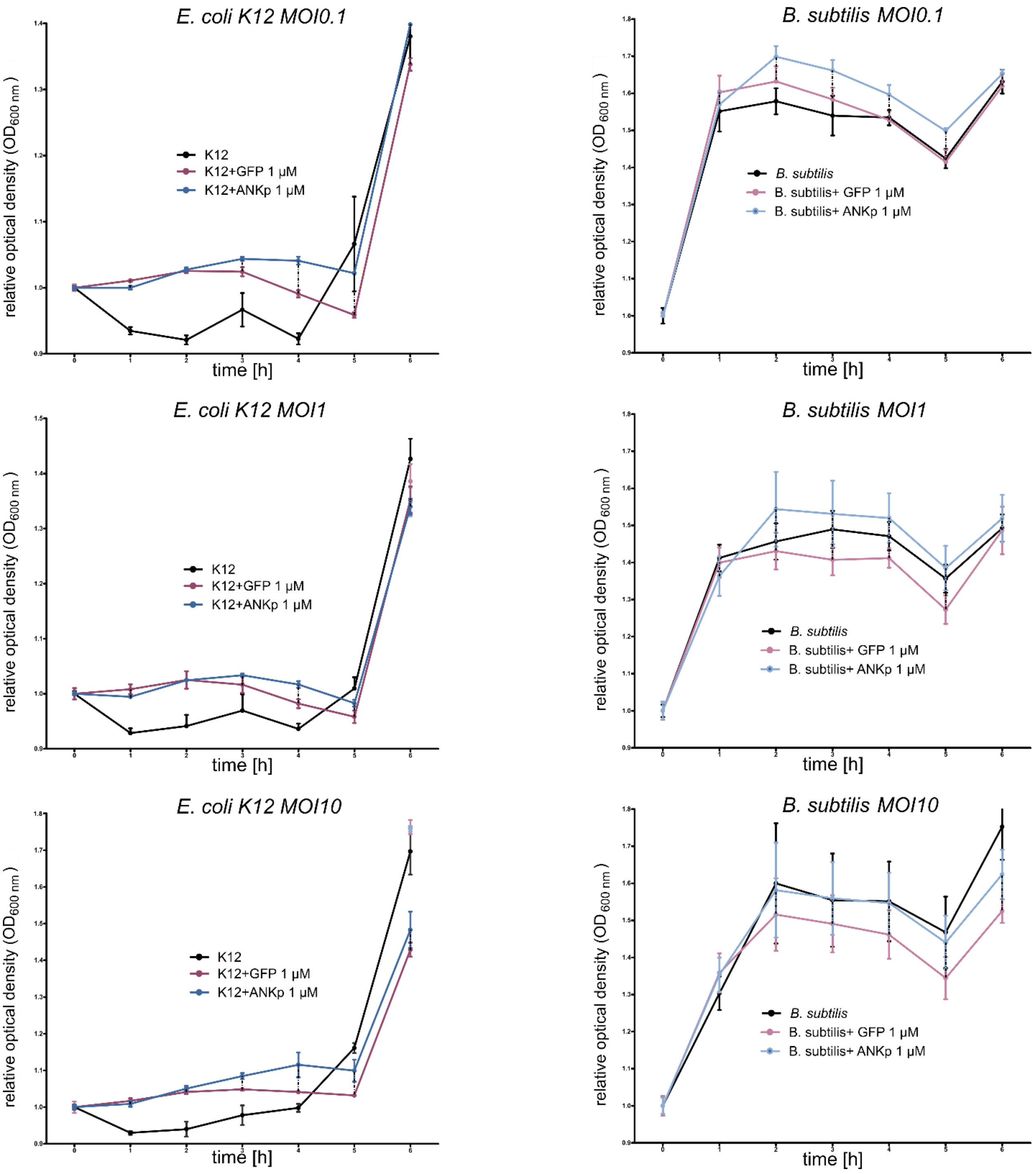


**Supplementary Figure 5: Growth kinetics of *E. coli* K12 and *B. subtilis* on macrophages (BMDMs) upon ANKp treatment.** Optical density (OD) was normalised to the start value at the beginning of the experiment ($t_0$). The multiplicity of infection (MOI) denotes the microbe:macrophage ratio set at $t_0$. Data are presented as the mean ± SEM from 4 independent replicates.

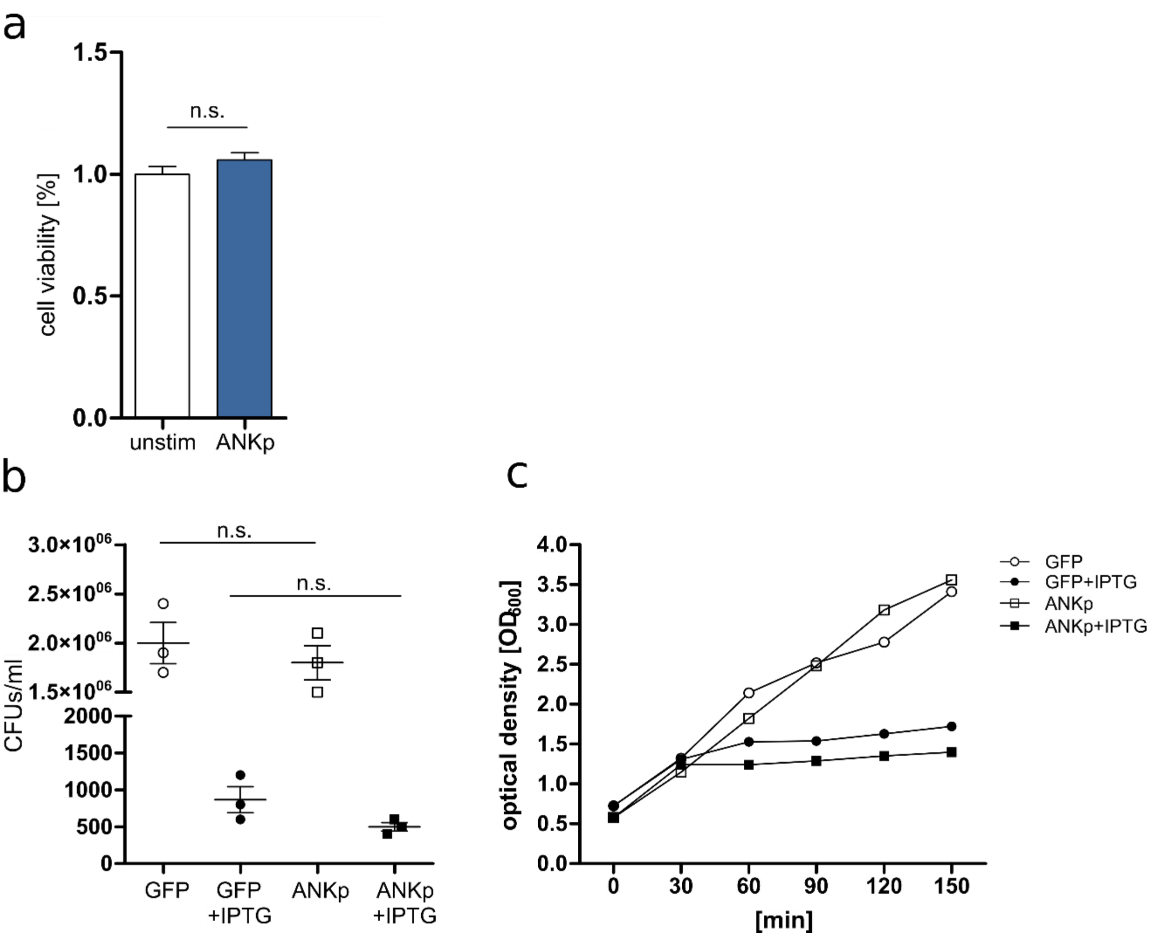


**Supplementary Figure 6: ANKp cytotoxicity testing.** (**a**) BMDM cell viability was measured with an MTS assay. Data are presented as the mean ± SEM from 4 replicates each. (**b**) *E. coli* viability on agar plates was measured by counting colony forming units with and without IPTG induction of ANKp expression. Data are presented as the mean ± SEM of 3 independent experiments. (**c**) *E. coli* viability in liquid LB-medium was measured by following optical density over time with and without IPTG induction of ANKp expression. Statistical significance between treatments was determined by two-tailed unpaired Student´s *t*-tests with $*p < 0.05$, $**p < 0.01$ and $***p < 0.001$.

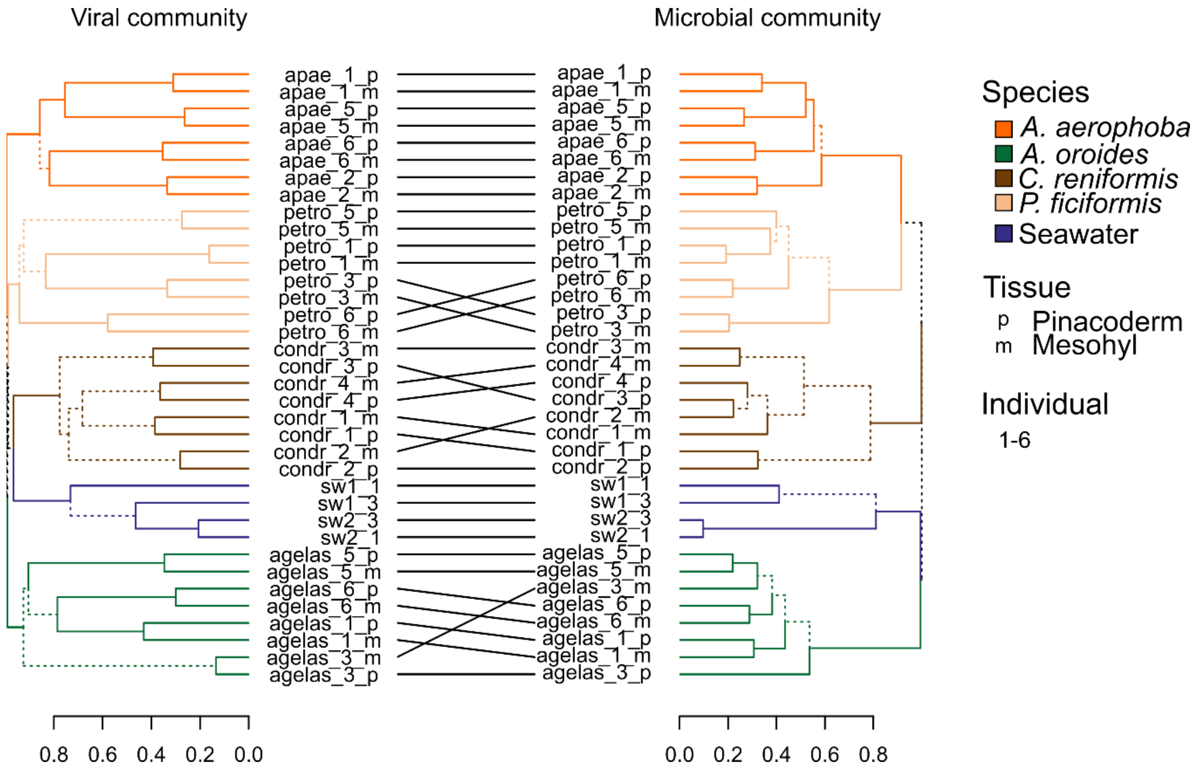


**Supplementary Figure 7: Hclust analysis supplementing CrAss and comparison to microbial community profiles.** Relative abundance profiles of (left) BCvir population contigs and (right) co-extracted prokaryotic 16S rRNA gene sequences, hierarchically clustered (complete linkage) with the Bray-Curtis similarity. Cladograms are compared with the tanglegram function (R dendextend package). Dashed lines indicate differences in tree topologies. Sample IDs indicate the following: apae, agelas, petro, condr are the sponge species *Aplysina aerophoba*, *Agelas oroides*, *Petrosia ficiformis*, *Chondrosia reniformis*. sw indicates seawater. _m and _p indicate mesohyl and pinacoderm tissues. _#_ is the individual identifier.

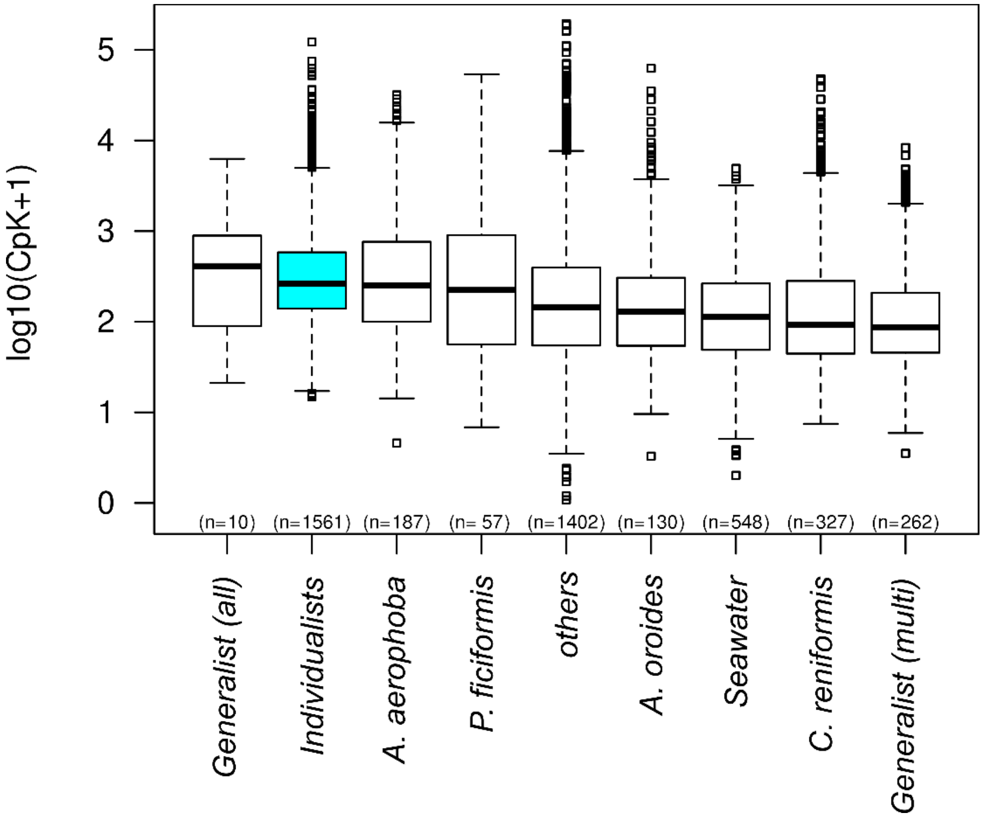


**Supplementary Figure 8: Relative abundance levels of BCvir prevalence groups.** The abundance estimate is counts per kilobase (CpK). The number of contigs per category is denoted by *n*. Samples were sorted by median abundance of prevalence classes.

### Supplementary Tables

**Supplementary Table 1.** Cross-Assembly statistics.

| **Criterium** | **Parameter** |
|---|---|
| Assembly tool(s) used | metaSPAdes; 3.11.0; default parameters |
| Number of assembled contigs (> 5000) | 7,105 |
| Total bases assembled (bp) | 72,419,424 |
| Contig N50 | 10,746 |
| Largest contig (bp) | 155,600 |
| % of Sequences assembled | 62.47% of the reads mapped back to whole cross assembly<br>51.41% of the reads mapped back to filtered viral contigs |

**Supplementary Table 2.** Kruskal–Wallis test followed by Dunn’s Post hoc-Tests with Benjamini-Hochberg false discovery rate correction.

| **pairwise comparisons** | **Z statistic (adjusted p value)** |
|---|---|
| BC_1608 - BC_2142 | 0.141592 (0.4437) |
| BC_1608 - BC_2160 | -0.161975 (0.4841) |
| BC_2142 - BC_2160 | -0.265500 (0.4941) |
| BC_1608 - neg_control | 3.665383 (0.0002)* |
| BC_2142 - neg_control | 3.324442 (0.0006)* |
| BC_2160 - neg_control | 3.350797 (0.0007)* |
| BC_1608 - pos_control | -7.922506 (0.0000)* |
| BC_2142 - pos_control | -7.421882 (0.0000)* |
| BC_2160 - pos_control | -6.666282 (0.0000)* |
| neg_control - pos_control | -9.349236 (0.0000)* |
| alpha = 0.05 | |
| Reject Ho if p <= alpha/2 | |

**Supplementary Table 3.** Sampling metadata.

| species | used for | location | date | time | temperature | depth | lat | long |
|---|---|---|---|---|---|---|---|---|
| *A. aerophoba* | virome | Port Lligat | 12. July 2016 | noon | 23.3°C | 3m | 42°17'50.8"N | 3°17'20.4"E |
| *P. ficiformis* | virome | Reserva Natural de Montgrí, Illes Medes i Baix Ter | 18. July 2016 | noon | 21°C | 15m | 42°04'56.97"N | 3°12'00.09"E |
| *C. reniformis* | virome | Reserva Natural de Montgrí, Illes Medes i Baix Ter | 18. July 2016 | noon | 21°C | 15m | 42°04'56.97"N | 3°12'00.09"E |
| *A. oroides* | virome | Reserva Natural de Montgrí, Illes Medes i Baix Ter | 18. July 2016 | noon | 21°C | 15m | 42°04'56.97"N | 3°12'00.09"E |

**Supplementary Table 4.** TaqMan assays used in this study.

| Gene | Company | TaqMan assay ID |
|---|---|---|
| Cxcl1 | Life Technologies | 433859 |
| Gapdh | Life Technologies | 99999915 |
| Ifnb1 | Life Technologies | 439552 |
| Tnfα | Life Technologies | 443258 |

### Supplementary References


1. Rodriguez RL, Gunturu S, Tiedje JM, Cole JR, Konstantinidis KT. Nonpareil 3: Fast Estimation of Metagenomic Coverage and Sequence Diversity. *mSystems* **3**,  (2018).

2. Brown JK*, et al.* Revision of Begomovirus taxonomy based on pairwise sequence comparisons. *Arch Virol* **160**, 1593-1619 (2015).

3. Laffy PW*, et al.* Reef invertebrate viromics: diversity, host specificity and functional capacity. *Environ Microbiol* **Epub ahead of print**,  (2018).

4. Nguyen LT, Schmidt HA, von Haeseler A, Minh BQ. IQ-TREE: a fast and effective stochastic algorithm for estimating maximum-likelihood phylogenies. *Mol Biol Evol* **32**, 268-274 (2015).

5. Sela I, Ashkenazy H, Katoh K, Pupko T. GUIDANCE2: accurate detection of unreliable alignment regions accounting for the uncertainty of multiple parameters. *Nucleic Acids Res* **43**, W7-14 (2015).
